# Modulation of glucose metabolism by 2-Deoxy-D-Glucose (2DG) promotes IL-17 producing human T cell subsets

**DOI:** 10.1101/2022.03.13.484135

**Authors:** Xin Chen, Lina Kozhaya, Chuxi Wang, Lindsey Placek, Ece Karhan, Derya Unutmaz

**Affiliations:** Jackson Laboratory for Genomic Medicine, Farmington, Connecticut, USA; Department of Immunology, University of Connecticut School of Medicine, Farmington, CT USA; Institute for Immunity, Transplantation, and Infection, Stanford University, Stanford, CA USA

## Abstract

Activation and differentiation of T cells are closely linked to their cellular metabolic programs. Glycolysis and mitochondrial metabolism are thought to be critical in modulating T cell function. Here we asked to what extent inhibition of glycolysis, using 2-Deoxy-D-Glucose (2DG), regulate activation, effector function, or differentiation of human T cell subsets. We found that glycolysis is required for T cell receptor (TCR) -mediated activation and proliferation of human naive CD4+ T cells but had less of an impact on memory subsets. CD4+ T cells cultured in the presence of 2DG displayed higher level of IL-17-secreting cells (Th17) from memory or *in vitro* differentiated naive regulatory T cell (Tregs) subsets. Moreover, the mucosal associated invariant T (MAIT) cell subset survived or expanded better and secreted higher IL-17 in the presence of 2DG. Remarkably, we found that the 2DG effect was reversed by mannose but not by glucose. Collectively, these findings suggest that 2DG could enrich IL-17 secreting human effector T cell subsets and their cellular functions. Our finding provides a framework to manipulate glycolytic pathways in human T cells in infectious diseases such as COVID19 and in enhancing cancer immunotherapy.

## Introduction

Metabolism has been recognized as an important regulator in T cell activation and lineage differentiation (Araki et al., 2010; Chapman et al., 2020; Jacobs et al., 2008; MacIver et al., 2013; Palmer et al., 2015; van der Windt & Pearce, 2012). Upon T cell activation, TCR signals and co-stimulation activate the phosphatidtyl-inositide-3 kinase (PI3K)/Akt/mTORC1 signaling pathway for the increased anabolic needs of effector T cells, which become more sensitive to metabolic regulation (Ho et al., 2015; Palmer et al., 2015; Sena et al., 2013). Importantly, an efficient immune response involves various T cell subsets, which have different metabolic requirements for development and effector functions (MacIver et al., 2013; Pearce et al., 2013). It is still unclear how metabolic pathways regulate the functions of T cell subsets specifically, but understanding the underlining mechanisms would allow modulation of immunity in chronic disease and cancer (Johnson et al., 2018).

Subsets of T cells that secrete IL-17 are the critical mediators in antimicrobial and pro-inflammatory responses, as well as in pathogenesis in autoimmune or chronic inflammatory diseases (Bettelli et al., 2006; Damasceno et al., 2020; Platt et al., 2020; Xu et al., 2020; Yasuda et al., 2019). Th17 cells and a subset of Tregs are the major sources of IL-17 secretion, which are also characterized by the expression of the transcription factor RORC and the chemokine receptor CCR6 (Singh et al., 2008; Valmori et al., 2010; Wan et al., 2011). Several mouse studies have revealed a role of glucose metabolism in Th17 differentiation, via mTORC1, Myc, and HIF1a signaling (Kastirr et al., 2015; Perl, 2016; Sasaki et al., 2016; Shen & Shi, 2019; Shi et al., 2011). Inhibition of glycolysis with 2-Deoxy-Glucose (2DG) or 3-bromopyruvate in mice for a short period of time impaired Th17 differentiation, phenocopying the effect of HIF1a deficiency (Okano et al., 2017; Shi et al., 2011). However, how glycolytic metabolism regulates the differentiation and functions of human Th17 and IL-17 secreting Tregs remains unclear.

Another source of IL-17 producing T cells are mucosal associated invariant T (MAIT) cells (Coulter et al., 2017; Willing et al., 2018). Human MAIT cells can be identified by a semi-invariant T cell receptor(TCR) alpha chain(Va7.2) and CD161 expression (Chiba et al., 2018). MAIT cells can be activated by a broad range of bacteria and yeasts, and can discriminate and fine-tune their functional responses to complex human microbiota (Constantinides et al., 2019; Corbett et al., 2020; Hinks & Zhang, 2020; Ioannidis et al., 2020; Tastan et al., 2018). Thus, IL17-secreting MAIT cells play a role in infections and autoimmune diseases such as Tuberculosis and Multiple Sclerosis (Coulter et al., 2017; Willing et al., 2018). While the frequency and distribution of MAIT cells are currently under intense investigation, there is still much to understand in the metabolic regulation of the function of the human MAIT cell subset.

In this study, we utilized 2DG, an analog and potent glycolysis inhibitor, to investigate the role of glucose metabolism in the activation, differentiation, and effector function of human T cell subsets. We showed that 2DG suppressed human T cell activation, cytokine production, and proliferation upon early T cell receptor (TCR) activation. However, 2DG treated naive T cells showed a greater reduced proliferative capacity and glycolytic function compared to memory T cells. Remarkably, 2DG treatment greatly enriched IL-17 producing CD4+ T cells in long term culture and during *in vitro* differentiation of naive or naive regulatory T cells (TNreg) or from the MAIT cell subset. Together, our findings suggest a differential impact of glycolysis in different subsets and differentiation stages of human T cells.

## Results

### Inhibitor of glucose uptake, 2DG, suppresses early activation of CD4+ T cells

It has been reported that 2DG inhibits glucose metabolism (Xi et al., 2014), which may in turn regulate T cell activation (Buck et al., 2015; Palmer et al., 2015). Therefore, we first examined the effect of 2DG on T cell activation by assessing the expression of the IL-2 receptor alpha (CD25) and the glucose transporter 1 (GLUT1), as both are upregulated upon T cell activation for IL-2 signaling and glucose uptake, respectively (Chapman et al., 2019). Human primary CD4+ T cells were activated with anti-CD3/CD28 beads in media alone or in media supplemented with 0.3mM, 1mM, or 3mM 2DG. The expression of CD25 and GLUT1 in T cells was significantly reduced by 3mM 2DG compared to the control (**Figure 1A-B**), but lower concentrations of 2DG (0.3mM or 1mM) did not reach statistical significance (data not shown). Because 2DG suppresses glycolysis (Xi et al., 2014), we also activated T cell subsets in the glucose-free media as a comparison. In contrast to the effect seen with 2DG, T cells activated in glucose-free-media showed higher GLUT1 expression and CD25 expression comparable to the control condition (**Figure 1A-B**). We also observed a reduced expression of program cell death 1 (PD1) and Lymphocyte Activating 3 (LAG3) on 2DG-treated T cells, which are known to be induced upon TCR activation (Lichtenegger et al., 2018) (**Figure 1C-D**). However, the expression of PD1 or LAG3 was not significantly different between T cells activated in glucose-free or regular media (**Figure 1C-D**).

**Figure 1.**
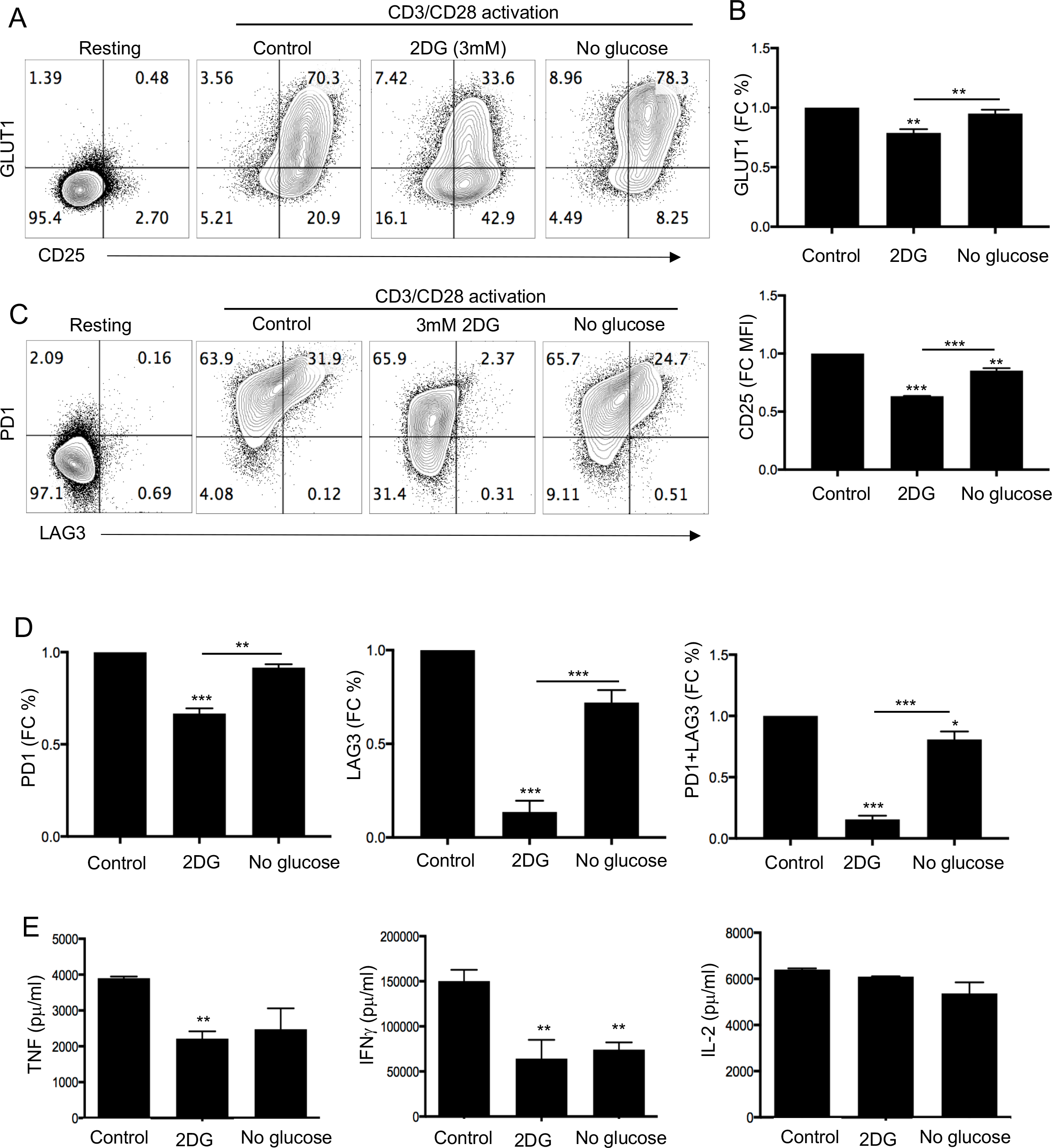
2DG suppresses early activation of CD4+ human T cells. T cell subsets were activated with anti-CD3/CD28 beads in media alone, 3mM 2DG or in glucose-free media. (A) Surface expression of CD25 and GLUT1 on resting and 24h-activated CD4+ T cells with indicated conditions. (B) Fold difference in GLUT1 expression and mean florescent intensity (MFI) of CD25 on 24h-activated CD4+ T cells in 3mM 2DG or in glucose-free media compared to control. (C) Flow cytometry plot of LAG3 and PD1 expression on resting and activated CD4+ T cell in media alone, 3mM 2DG, or in glucose-free media for 24h. (D) Statistical analysis of fold difference in PD1, LAG3, and double positive expression of CD4+ T cells activated for 24 h in 3mM 2DG and in glucose-free media. (E) CD4+ T cells were activated with anti-CD3/CD28 beads. Cell supernatant were collected from the indicated conditions after 2 days post beads activation. The levels of TNF, IFNg, and IL-2 cytokines were determined by FlowCytomix Multiplex bead assay. Data represent three independent experiments. **p <0.01, ***p <0.001

TCR-activation of T cells leads to a rapid production of cytokines such as TNF, IFNγ, and IL-2. As such, to determine whether 2DG also suppresses cytokine production, we collected activated T cell supernatants 2 days post CD3/CD28 activation and analyzed the cytokine levels using a cytometric bead assay. We found that the levels of TNF and IFNγ secretion were significantly reduced but there was no difference in IL-2 secretion in the presence of 2DG culture compared to control media (**Figure 1E**). Together, these data suggest that 2DG suppresses early human T cell activation and cytokine production.

### 2DG has different effects on activation or effector functions of naive and memory T cells

It has been reported that naive and memory T cells may have different metabolic requirements for activation and proliferation (Almeida et al., 2016; MacIver et al., 2013; van der Windt & Pearce, 2012). To determine whether 2DG modulates the activation of naive vs memory CD4+ T cells differently, we sorted these subsets from human CD4+ T cells using CCR7+CD45RO- (naive) and CCR7+/-CD45RO+ (memory) markers and activated with anti-CD3/CD28 beads in media alone, in 3mM 2DG, or in glucose-free media. Both T cell subsets displayed a similar trend of reduction in GLUT1 and CD25 expression on day 1 post activation **(Figure 2A)**; however, on day 4 post-activation, 2DG-treated memory T cells upregulated GLUT1 to levels comparable to control cells whereas most naive T cells remained GLUT1 negative **(Figure 2B-C)**. Similar to unsorted CD4+ T cells (**Figure 1A**), activation of naive or memory T cells in glucose-free media did not significantly affect the expression of GLUT1 or CD25 (**Figure 2A-C**). Addition of IL-2 in these activation experiments did not restore the 2DG-mediated down-regulation of GLUT1 or CD25 expression in either naive or memory T cells (**Supplemental Figure 1A-C**). Since we have observed that 2DG suppresses T cell activation and glucose uptake, we hypothesized that 2DG would also reduce T cell proliferative capacity. To address this, we labeled naive and memory T cells with Celltrace violet (CTV) dye, and then activated with anti-CD3/CD28 coated beads. Indeed, in 2DG-treated or in glucose-free media, naive or memory T cell subsets proliferated less compared to control media (**Figure 2E**). Addition of IL-2 did not restore the low proliferation of 2DG-treated T cell subsets (**Supplemental Figure 1D**).

**Figure 2.**
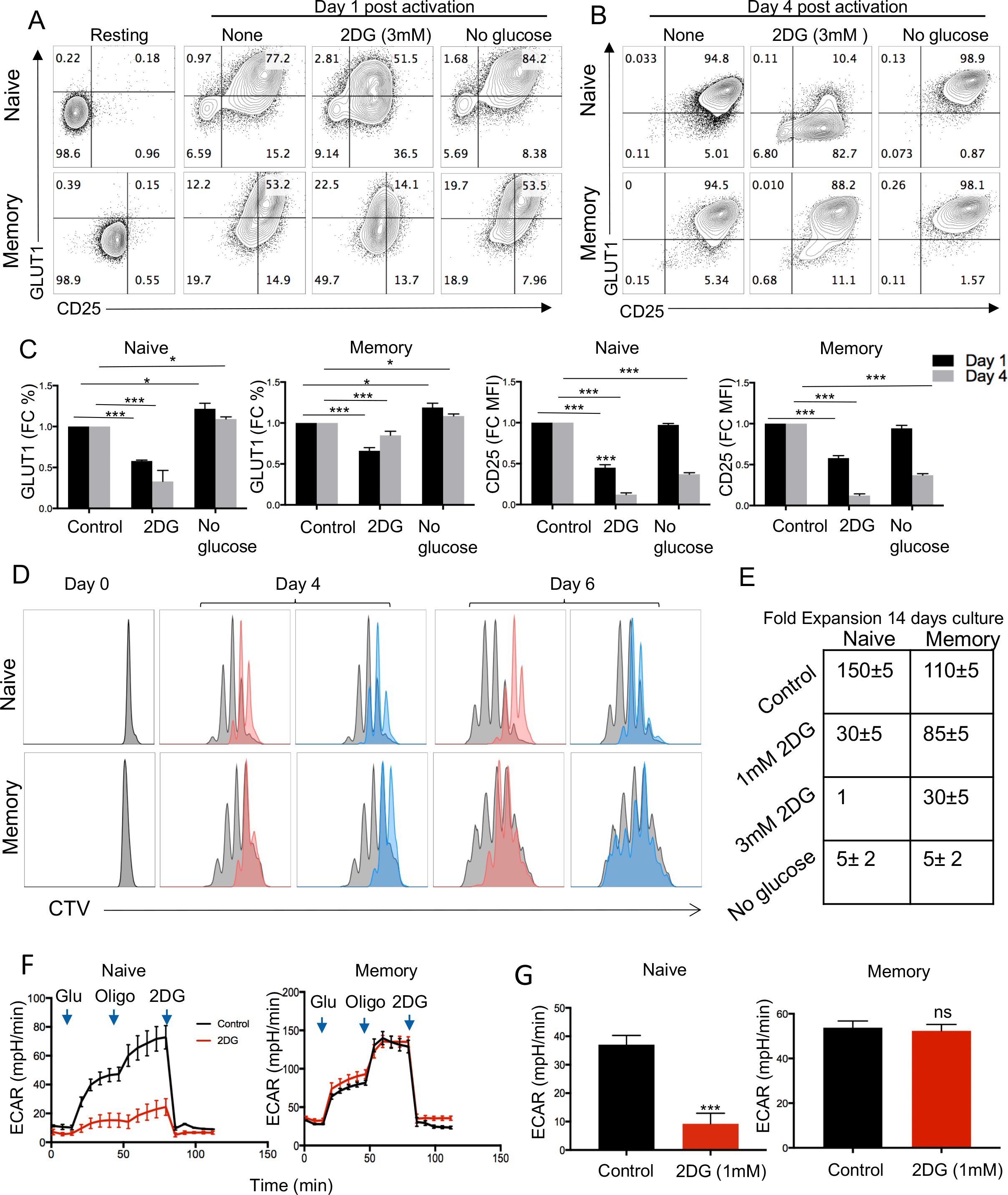
2DG has differential effects on activation of naive and memory T cell subsets. (A) Naive and memory CD4+ T cells were activated with anti-CD3/CD28 beads in media alone, 3mM 2DG, or in glucose-free media. Media was not supplemented with IL-2. Representative flow cytometry plot of GLUT1 and CD25 surface expression on resting, and day 1(A)- or day 4(B)- post activated Naive (top) and memory T cells (bottom) in indicated conditions were shown. (C) Fold difference in frequency of GLUT1 expression or mean fluorescence intensity of CD25 expression on naive or memory T cells on day 1 (black)- and day 4 (grey)-post activation in indicated conditions. (D) Naive and memory CD4+ T cells were labeled with CellTrace violet (CTV) dye followed by anti-CD3/CD28 beads activation in indicated conditions. Representative histogram plot of CTV-labeled naive (up) and memory (bottom) CD4+ T cells in 3mM 2DG (red) or in glucose-free media (blue) after day 4 and day 6 post TCR activation. (E) Representative fold expansion of naive and memory CD4+ T cells in 1 or 3mM 2DG or in glucose-free media after 2 week T cells expansion. (F and G) Naive and memory CD4+ T cells were activated and expanded in media alone (control, black) or 1mM 2DG (red) for 14 days. Metabolic functions were analyzed using Seahorse glucose stress test assay. The extracellular acidification rate (ECAR) was assessed after the addition of glucose (gluc), oligomycin (oligo), and 2DG at indicated times and the glycolytic capacity was determined. Each set of data is representative of three donors. *p < 0.05, **p < 0.01, ***p<0.001

We next monitored the expansion of these activated T cells during 2-week culture in IL-2. Expansion of 1 mM 2DG treated naive T cells were 30 fold after 14 days, compared to 150 fold in the control, whereas memory T cells in the presence of 1mM 2DG expanded 85 fold compared to 110 fold in the control media. We then asked whether 2DG has differential effects in the glycolytic function of naive and memory subsets by using the Seahorse glucose stress test assay. After T cell expansion in 2DG or control media for 14 days, 2DG was washed away and cells were resuspended in Seahorse XF base media, and the metabolic functions were analyzed by the Seahorse XFe96 Analyzer. The extracellular acidification rate (ECAR) was assessed after the addition of glucose (gluc), oligomycin (oligo), and 2DG at indicated times and the glycolytic capacity was determined. 2DG-treated memory T cells had comparable glycolytic function to the control groups; however, 2DG-treated naive T cells showed significantly reduced glycolysis in response to addition of glucose (**Figure 2F-G**).

### 2DG enhances IL-17 producing CD4+ memory subsets

We next further explored which effector T cell subsets are modulated by suppressing glycolysis. We first observed a significantly increased CD161+ and CCR6+ populations in 2DG-treated CD4+ T cells compared to control (**Figure 3A and 3B**). Since nearly all IL-17 secreting cells express CCR6 and CCR6+ Th17 cells are reported enriched in the CD161+ subset (Acosta-Rodriguez et al., 2007; Cosmi et al., 2008; Wan et al., 2011), we hypothesized that 2DG would preferentially affect the function of IL-17 secreting T cell subsets. Indeed, after two weeks in culture with 2DG, we observed that IL-17+ subsets were increased in both CCR6+ and CD161+ populations compared to cells in control media, but interestingly, there was no difference in the low glucose condition (**Figure 3C**). After 2DG was removed from the culture and cells reactivated with phorbol myristate acetate (PMA) and calcium ionophore (ionomycin), we determined intracellular cytokine secretion in the presence of brefeldin a (GolgiStop). We found that the frequency of cells that expressed not only IL-17 but also IFNγ, IL-4, and IL-21 were significantly increased (**Figure 3C and 3D**). However, no effect was seen in IL-2 and TNF production from CD4+ T cells (**Supplemental Figure 3**). Taken together, these data demonstrate that 2DG-treatment can modify the cytokine production of lineage-committed cells in long term *in vitro* cultures.

**Figure 3.**
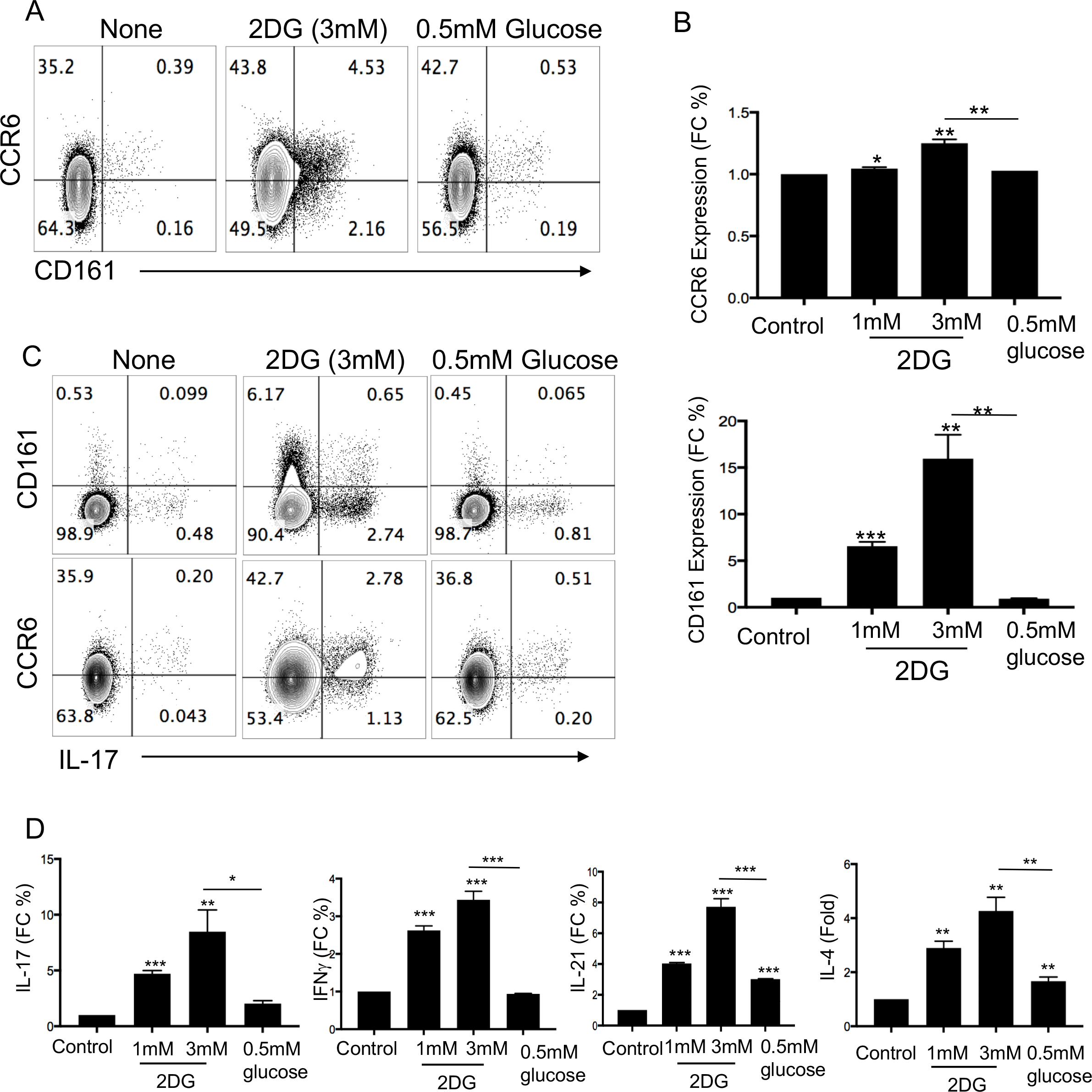
2DG enhances IL-17 production in primary CD4+ T cells. Purified CD4+ T cells were activated with aCD3/CD28 beads in media alone, 3mM 2DG, and in media with 0.5mM glucose. (A) The representative flow cytometry plot of CCR6 and CD161 expression on day14 post- activated T cells were shown. (B) Statistical analysis of fold difference in frequency of CCR6+ and CD161+ T cells in indicated conditions. (C) After 14 days’ cell culture, T cells activated and expanded in indicated condition were re-stimulated with PMA/ionomycin for 4h, followed by intracellular staining as described in the method section. The percentage of IL-17 production from CD161+ and CCR6+ T cells were shown. (D) Statistical analysis of fold difference in frequency of IL17+, IFNg, IL-4, and IL-21+ T cells from indicated conditions was shown. Each set of data is representative of three donors. *p < 0.05, **p < 0.01, ***p<0.001

### 2DG enriches IL-17+ cells within Th17 lineage-committed CCR6+ memory T cell subset

Since we observed a significant increase in IL-17 cytokine production from 2DG-treated T cells, we hypothesized that 2DG could enhance the IL-17 production from Th17 cells compared with other effector T cells. To test this hypothesis, we sorted CD4+ T cells into CCR6- and CCR6+ memory T cell subsets, activated them in the presence or absence of 2DG, and expanded for two weeks in IL-2, as described above (**Figure 3A**). Consistent with our prior results, 2DG-treated CCR6- and CCR6+ CD4+ T cells displayed higher CD161 expression than control cells (**Figure 4A**). IL-17 production from purified and expanded CCR6+ T cells were increased in the presence of 2DG (**Figure 4B**). Notably, IFNγ+ and IL-21+ producing CCR6+ T cells were also increased by 2DG (**Figure 4B-E**). Interestingly, the percentage of IL-10+ and IL-17+ T cells was significantly increased in 2DG-treated CCR6+ T cells (**Figure 4F and 4G)**. Therefore, we concluded that 2DG increased CD161+ within both CCR6+ and CCR6- T cells, and also enhanced IL-17, IFNγ, IL-21, and IL-10 production from the sorted CCR6+ memory T cell subset.

**Figure 4:**
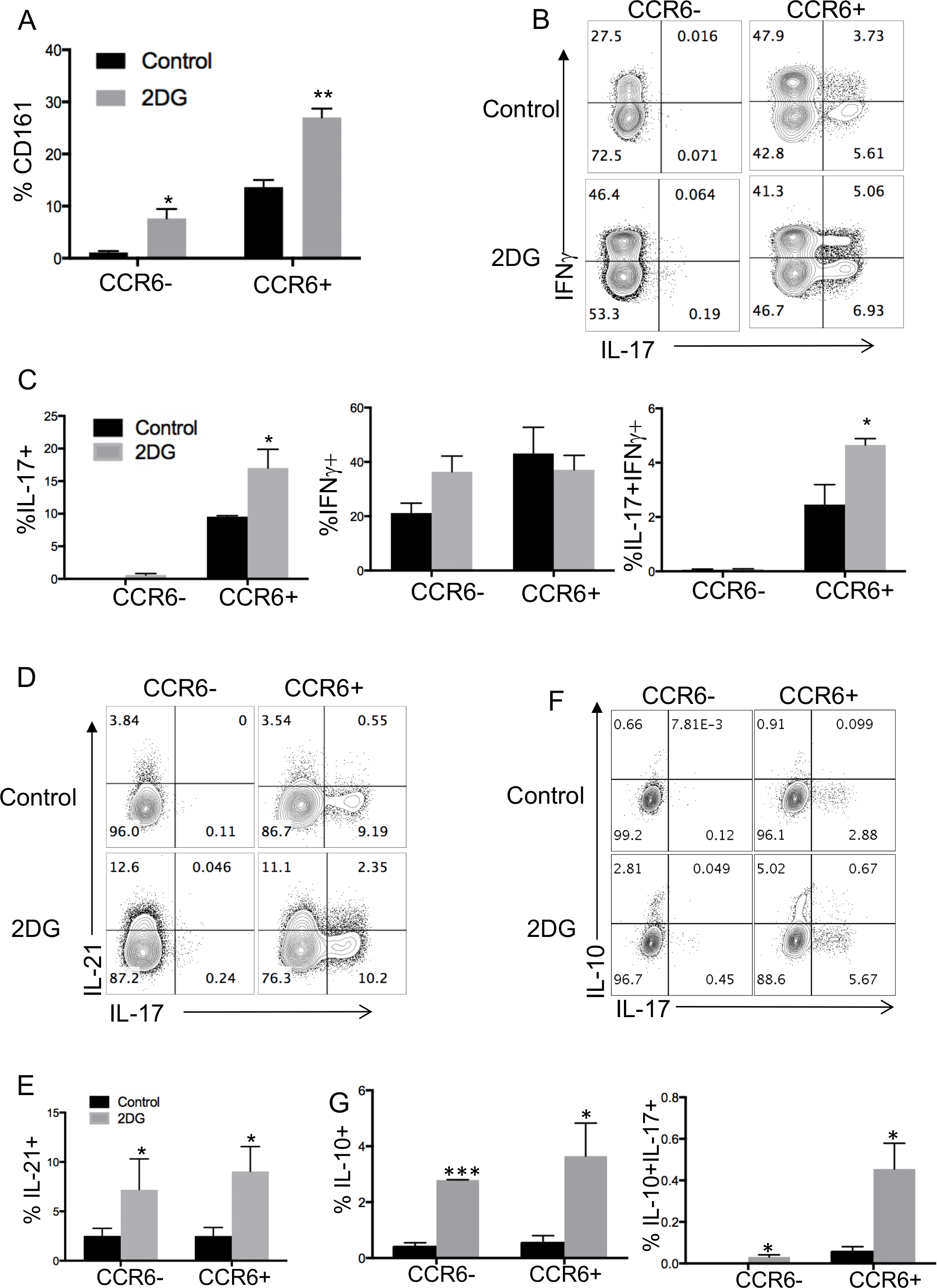
2DG enriches the frequency of IL-17 from sorted CCR6+ memory T cells. (A) sorted CCR6+ and CCR6- memory cells were activated with aCD3/CD28 beads in media alone or with 3mM 2DG for 14 days. The frequency of CD161+ cells in day-14 post activated sorted CCR6+ and CCR6- subsets in 3mM 2DG compared to control were shown. (B) 14 days post activated T cells were re-stimulated with PMA and ionomycin for 4h, followed by surface and intracellular staining. The frequency of IFNg and IL-17 was determined from sorted CCR6- and CCR6+ T cells. (C) Statistic analysis of the frequency of IL-17+, IFNg +, and IL17+ IFNg+ T cells from CCR6- and CCR6+ T cells as in Figure 4B. The representative plot (D) and the statistical analysis (E) of the frequency of IL-21, IL-10, and IL-17 was determined from sorted CCR6- and CCR6+ T cells after 14 days expansion and re-activation with PMA and ionomycin. The representative plot (F) and the statistical analysis (G) of the frequency of IL-10 and IL-17 was determined from sorted CCR6- and CCR6+ T cells after 14 days expansion as in Figure 4D and E. Each set of data is the representative of three donors. *p < 0.05, ***p < 0.001

### 2DG increases in vitro generation of IL-17-producing Tregs from human naive regulatory T cell (TNreg) precursors

A subset of Tregs that secretes IL-17 can preferentially arise from human naive precursors (CD25+CD45RO-CCR7+ cells, termed TNreg) in the presence of polarizing-cytokines IL-1β, IL- 23, and TGF-β (Mercer et al., 2014; Valmori et al., 2010). To explore how 2DG regulates Th17 cell differentiation from TNregs, we used this cytokine polarization protocol (**Figure 5A**) to differentiate and expand IL-17-secreting cells from highly purified naive or precursors naive regulatory T cells (TNregs) in media alone or in the presence of 1 mM 2DG. The FOXP3+ cells during polarization of TNregs with 2DG were statistically similar in frequency (**Figure 5B**). However, 2DG significantly increased generation of IL-17+ Tregs from TNreg precursors (**Figure 5C**). Since HELIOS expression defines a phenotypically distinct population of Tregs (Thornton et al., 2019), and *in vitro*-generated IL-17+ cells from TNreg precursors did not express HELIOS (Mercer et al., 2014), we then asked whether 2DG could induce IL-17, IFNγ, and IL-21 production from HELIOS+ or HELIOS- populations of naive and TNregs. We also compared the 2DG effects between HELIOS+ and HELIOS- population of naive cells since HELIOS plays a role in naive T cell differentiation (Ng et al., 2019). The frequency of IL-17+ T cells witihn HELIOS- naive T cells was increased by 2DG in polarizing condition, but not within HELIOS+ subset (**Supplemental Figure 4**). 2DG also significantly increased IL-17 and IFNγ populations from HELIOS- population from TNregs in the polarization condition (**Figure 5D-F**). Notably, when comparing the intracellular IL-21 from HELIOS+ and HELIOS- subset of Tregs, the frequency of IL-21+ cells was significantly higher in 2DG-treated HELIOS- subset (**Figure 5G and 5H**). But there was no significant difference in IL-21 cytokines secretion in HELIOS+ or HELIOS- Treg subsets in 2DG-treated naive T cells compared to controls (**Figure 5H**). Intracellular IL-10 was also increased in HELIOS- population of TNregs in polarizing cytokines, and was significantly increased in HELIOS+ and HELIOS- population from naive T cells (**Figure 5J**). In addition, increased cytokine production of IL-17A, IL-17F, IL-10, and IFNγ was also confirmed by corresponding levels of cytokines detected in the culture supernatants (**Figure 5K**). Taken together, 2DG enhanced the generation of IL-17- producing cells from both naive and TNregs, which was restricted to HELIOS- subsets. 2DG also increased the production of IFNγ, IL-21, and IL-10 from TNregs in the polarizing condition.

**Figure 5.**
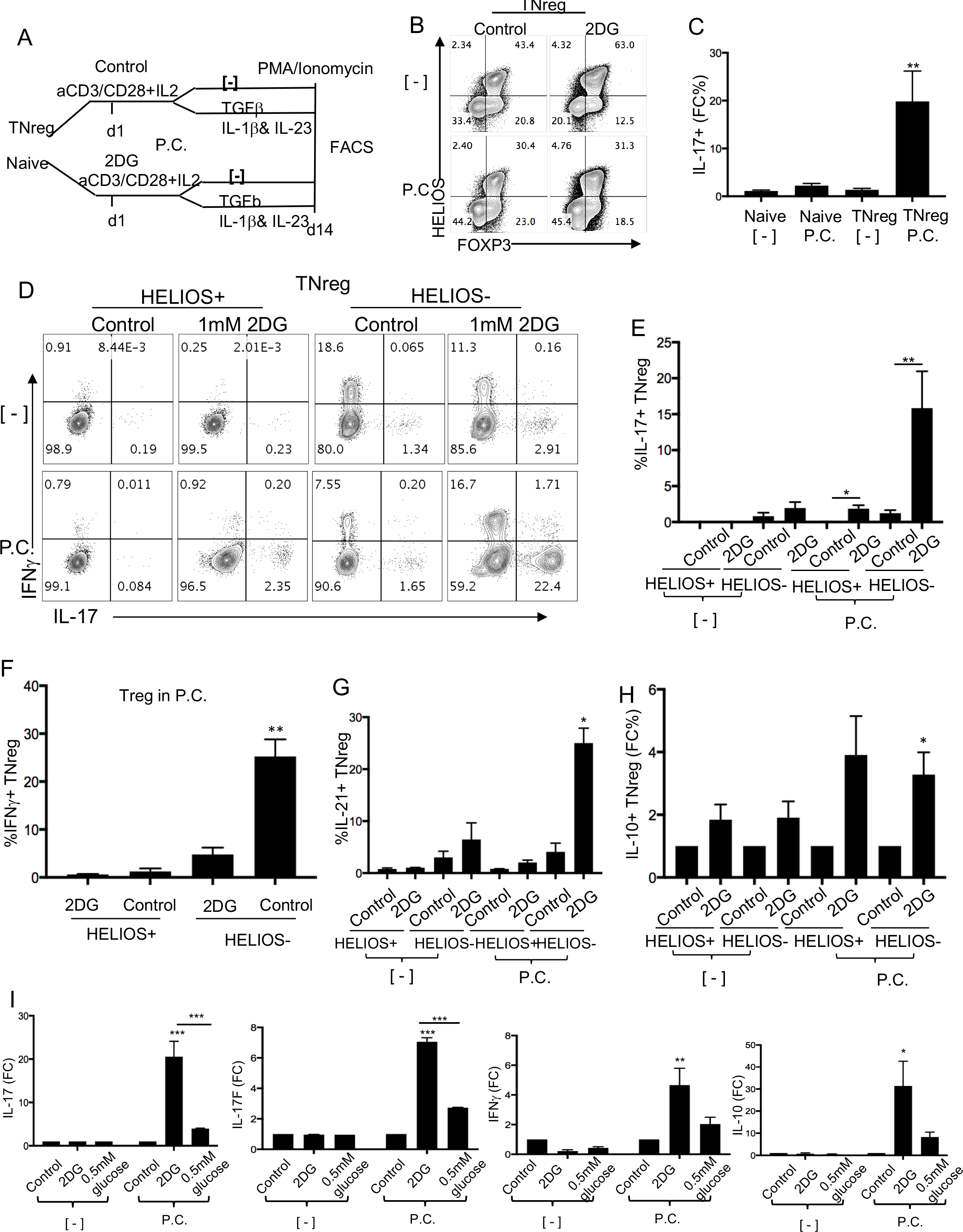
2DG enhances *in vitro* generation of IL-17-producing cells from TNreg cells. (A) Polarization protocol with Naive or TNreg cells. (B) Representative FOXP3/HELIOS expression of TNreg in two-week polarization cultures with or without 1mM 2DG. (C) Fold difference of IL-17 production from 1mM 2DG treated TNreg cells with polarizing cytokines (P.C.) or without (-) compared with control after 14-day expansion. (D) Representative flow cytometry plots of intracellular IFNγ and IL-10 secretion within HELIOS+ and HELIOS- gated populations of TNreg after 14-day expansion. (E) Statistical analysis of IL-17 production of indicated subsets from 1mM treated naive T cell (left)/TNreg(right) with or without polarizing cytokines. (F) Statistical analysis of IFNγ from 1mM treated TNreg in polarizing cytokines (G) Representative flow cytometry plots of intracellular IL-21 and IL-17 secretion within HELIOS+ and HELIOS- gated populations of TNreg after 14-day expansion. (H) Statistical analysis of IL-21 production of indicated subsets from 1mM treated naive T cell (left)/TNreg(right) with or without polarizing cytokine s. (I) Representative flow cytometry plots of intracellular IL-10 and IL-17 secretion within HELIOS+ and HELIOS- gated populations of TNreg after 14-day expansion. (J) Statistical analysis of fold difference of IL-10 production of indicated subsets from 1mM treated naïve T cell (left)/TNreg(right) with or without polarizing cytokines. (K) Day-14 expanded TNreg with or without polarization condition were re-stimulated with anti-CD3/CD28 beads. Cell supernatant were collected from the indicated conditions after 2 days post beads activation. IL-17A, IL-17F, IFNγ, and IL-10 productions from supernatant were determined by FlowCytomix Multiplex bead assay. Data represent three independent experiments. *p < 0.05, **p < 0.01, ***p<0.01

### Mannose reverses or rescues 2DG effects on T cell activation and effector functions

2DG has previously been shown to inhibit *N*-linked glycosylation (Andresen et al., 2012; Xi et al., 2014), which can be modulated through a branch of the glucose metabolism pathway called the hexosamine biosynthetic pathway (HBP) (Akella et al., 2019). Further, mannose, which is a major component of sugar moeity in glycoproteins, can reverse the apoptotic effect of 2DG on cancer cells (Ahadova et al., 2015; Gu et al., 2017; Kurtoglu et al., 2007). Thus, we sought to determine whether mannose can also reverse the 2DG effects on short term activation of T cells (**Figure 2**). Accordingly, we activated CD4+ naive T cells with anti-CD3/CD28 beads in media alone, with 3mM 2DG or mannose alone, or with 3mM 2DG supplemented with 3mM mannose. While we did not observe any difference in CD25 and GLUT1 expression on day 1 post activation (**Supplemental Figure 5**), mannose reversed the 2DG-mediated downregulation of CD25 and GLUT1 expression on day 4 (**Figure 6A and 6B**). Additional equimolar glucose did not reverse the 2DG effects (**Figure 6B**). Equal molar of mannose significantly reverses the proliferative capacity from 1 fold expansion with 2DG alone to 50 fold expansion with 2DG and mannose comparing to 157 fold cell expansion in control media (**Figure 6C**). However, after long-term culture, mannose did not rescue the impaired glycolytic function of 2DG-treated naive T cells (**Figure 6D and 6E**).

**Figure 6.**
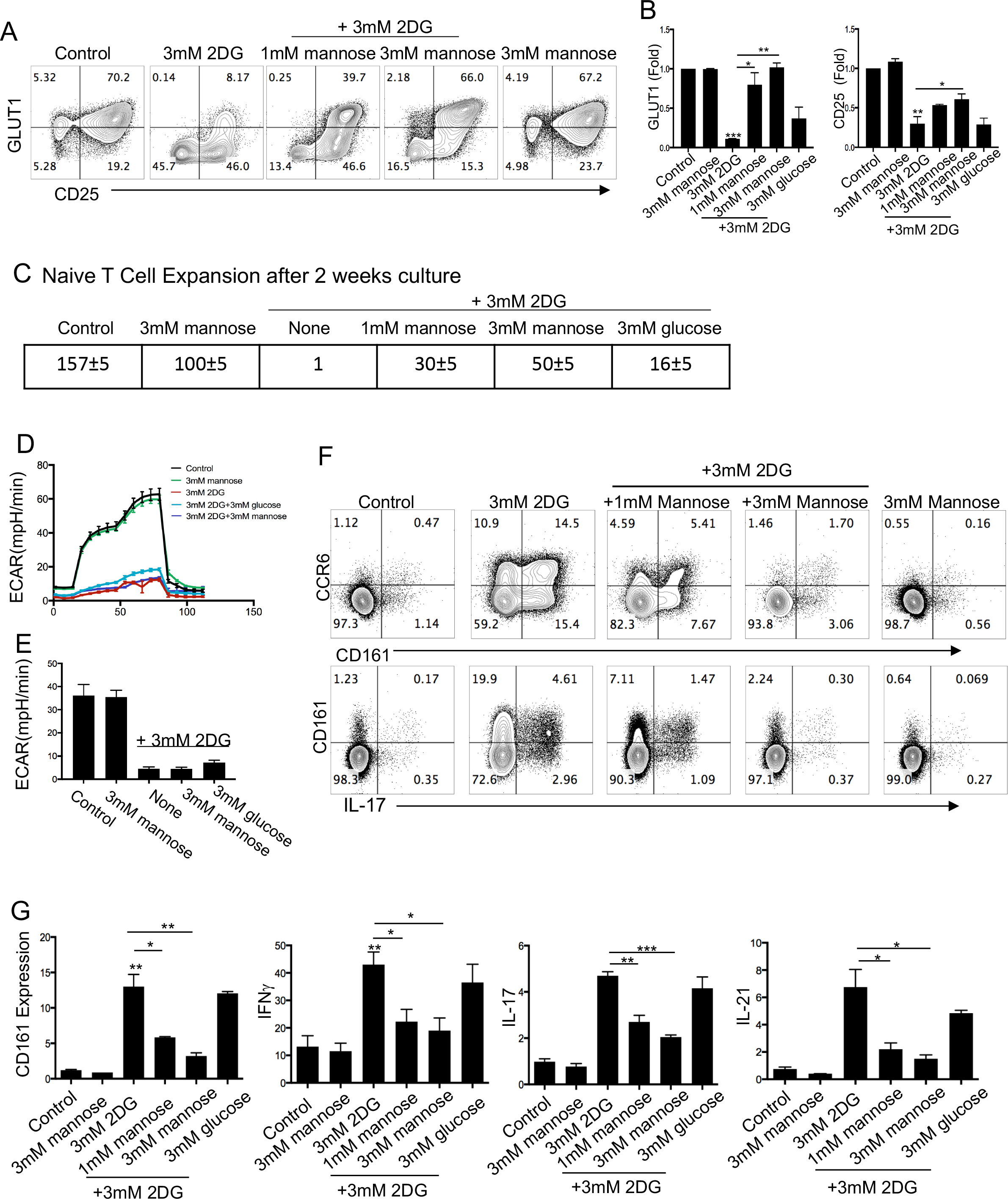
Mannose reverses or rescues 2DG effects on T cell subsets. (A) Naive CD4+ T cells were activated with aCD3/CD28 beads in indicated conditions. Representative flow cytometry plots of GLUT1 and CD25 surface expression. (B) Statistical analysis of fold difference in GLUT1 and Mean Florescent Intensity (MFI) of CD25 on activated naive T cells as in Figure 6A. (C) Representative fold expansion of naive CD4+ T cells in indicated conditions after 2-week T cells expansion. (D and E) Naive T cells were activated and expanded in the indicated conditions for 14 days. 2DG was washed away and the metabolic functions were analyzed using Seahorse glucose stress test assay. The extracellular acidification rate (ECAR) was assessed after the addition of glucose (gluc), oligomycin (oligo), and 2DG at indicated times and the glycolytic capacity was determined. (F) CD4+ T cells were activated with aCD3/CD28 beads in media alone, 3mM 2DG alone, 3mM 2DG plus 1mM or 3mM mannose. 14-days post activated T cells were re-stimulated with PMA and ionomycin for 4h, followed by surface and intracellular staining. Representative flow cytometry plots and statistical analysis of T cells expressed CCR6, CD161, IL-21, and IL-17. Data represent three independent experiments. *p < 0.05, **p < 0.01, ***p<0.01

Next, we asked whether mannose could reverse 2DG effects on IL-17 enrichment. After 14 days cell expansion, we observed a reduction of CCR6+ and CD161+ population in a mannose-does dependent manner (**Figure 6F and 6G**). In addition, the 2DG mediated-reduction of IL-17, IFNγ, and IL-21 were also reversed by mannose (**Figure 6G**). Addition of equimolar glucose into 2DG culture did not reverse the 2DG-mediated enrichment of IL-17 production (**Figure 6F and 6G**). Together these findings suggest that the inhibitory effects of T cells activation and IL-17 production on 2DG-treated human T cells are mannose-dependent.

## Discussion

In this study, we demonstrated that the glucose analog 2DG have remarkably different effects on activation and differentiation of human T cell subsets. 2DG suppresses early human T cell activation through the TCR, as assessed by cytokine production, and proliferation. However, in long term cultures 2DG also profoundly increases the frequency of IL-17 production from lineage-committed memory CCR6+CD4+ (Th17) T cells, MAIT cells and Tregs. In addition, inhibition of glycolysis by 2DG in these memory subsets (Th17, MAIT and Tregs) does not affect their expansion upon activation. On the other hand, naive CD4+ T cells are more sensitive to the inhibition of glycolysis by 2DG and therefore have reduced their survival and function. Importantly, we also demonstrated that the 2DG-mediated IL-17 enrichment could be reversed by addition of equimolar mannose.

A major question derived from our findings is that how inhibition of glycolysis can significantly increase the IL-17 production from already lineage-committed/differentiated memory Th17 cells? Several mice studies have revealed a role of glucose metabolism in Th17 differentiation (Kastirr et al., 2015; Perl, 2016; Sasaki et al., 2016; Shen & Shi, 2019; Shi et al., 2011) but showed discrepancies on the effect of inhibiting glycolysis in Th17 differentiation and functions. For example, in one study Tregs treated with 0.2mM 2DG for 5 days reduced IL-17 production *in vitro* (Li et al., 2019) and in another study reduced production of IL-17 by T cells after 5 day-2DG treatment was observed (Shi et al., 2011). However, another study reported an enhancement of Th17 cell polarization and IL-17 production after treatment of 2mM 2DG (Brucklacher-Waldert et al., 2017). In our study, we observed reduced IL-17 production by 2DG treated T cells for during early activation period. However, frequency of IL-17 producing subset was greately increased after long-term T cell culture (12-14 days), during reactivation in the absence of 2DG. Th17 can be characterized as “pathogenic” and “non-pathogenic” Th17 cells (Wu et al., 2018). Pathogenic Th17 cells express more effector molecules such as CXCL3, CCL4, CCL5, IL-3, and IL-22, whereas non-pathogenic Th17 cells exhibit upregulation of immune suppressive molecules and cytokines such as IL-10 (Gaublomme et al., 2015; Lee et al., 2012). We hypothesized that the increased subset of IL-17-secreting Th17 cells belongs to the “non-pathogenic” subset, as we also found an increase in IL-10 production from the same cells. Since IL-17+ Th17 cells are crucial for mediating mucosal immunity against fungi and bacteria infection in human subjects (Brembilla et al., 2018; McDonald, 2012), our findings may suggest that the metabolic changes affect the function of Th17 against commensal microbes in the gastrointestinal tract which would lead to mucosal damage and mucosal-relative diseases such as inflammatory bowel disease (IBD) (Galvez, 2014). Since the functions of Th17 cells in the small intestine can be closely regulated by gut microbiome (Evans-Marin et al., 2018; Garidou et al., 2015), our findings also suggest that a dysregulated metabolites from dysbiotic ileum microbiota (Visconti et al., 2019) could impact Th17 homeostasis, and were sufficient to induce metabolic disease such as Type 2 Diabetes (T2D) (Garidou et al., 2015).

We found that 2DG also increased the frequency of IL-17+ Tregs and IL-17 cytokine secretion from these cells in the presence of polarizing cytokines *in vitro* differentiation. These IL-17- producing Tregs were found to be accumulated in the inflamed joints of patients with rheumatoid arthritis and were functionally suppressive (Afzali et al., 2013; Jung et al., 2017). However, we did not observe a significant difference in suppressive capacity of 2DG-treated Tregs in proliferation assay (data not shown), which is consistent with a human Tregs study *in vitro* (Tanimine et al., 2019). Inhibition of glycolysis may also contribute to mucosal diseases, as IL-17+Tregs may contribute to the development of colon cancer (Knochelmann et al., 2018; Marshall et al., 2016), and possibly to IBD (Galvez, 2014).

As noted, an increased level of IL-10 from memory T cells (such as Tregs in the presence of polarizing cytokines, CCR6- and CCR6+ memory T cells) was also observed throughout the experiments. IL-10 is an anti-inflammatory cytokines, plays a central role in limiting the regulation of immune responses (Iyer & Cheng, 2012). Elevated IL-10 signaling can inhibit antigen presenting cells maturation, and chemokine secretion of the host during chronic viral infection (Granelli-Piperno et al., 2004) and autoimmune disease (Braat et al., 2003). It can be highly produced by a subset of Tregs, called Type 1 regulatory cells (Tr1) (Zeng et al., 2015), which is associated with autoimmune diseases such as IBD, multiple sclerosis(MS), and type 1 diabetes mellitus when their frequency was found reduced (Jia et al., 2019). Thus our data may suggest a potential role of glycolysis in the function of IL-10 secreting subsets and related diseases.

Another subset of cells, where found differential and specific effect of 2DG were MAIT cells, which are characterized with expression of CD161 and a semi-invariant Va7.2+TCR, are activated by a ligand from riboflavin metabolism called 5-ARU in the context of MR-1 molecule (Godfrey et al., 2019). MAIT cells are also unique in that they can be CD8+CD4- or CD8-CD4- and can also produce IL-17 and mostly migrate to mucosal regions, thus potentially play an important role in mucosal homeostasis and microbiome regulation (Oh & Unutmaz, 2019). In our experiments, we had found that 2DG-treated cells compared to control induced higher frequency of CD161+ cells, which is also highly expressed on MAIT cells (CD161hiVa7.2+) (Cosmi et al., 2008; Kleinschek et al., 2009). We also identified a distinct subset exhibiting medium-expression of CD161 which is also enhanced by 2DG, but is neither Va7.2+ (MAIT) nor CCR6+ (Th17) cells. The identity of this subset is not fully clear, but Klenerman et al reported a distinct functional subset in human T cells which had mid-level CD161 expression, produced IL-17/IFNγ, and were found to be highly enriched in chronic inflammation (Billerbeck et al., 2010). Future studies will be needed to better characterize this subset and its functions, as these have also shown different glucose tolerance than other effector cells.

Our findings that 2DG enhances IL-17 production also in MAIT cells, in addition to Th17 cells, suggest a shared metabolic pathways that regulate the effector functions of these mucosa-associated subsets. Interestingly IL-17+ MAIT cells are found specifically enriched from the peripheral blood of multiple sclerosis (MS) patient, implicating a proinflammatory role in autoimmune diseases (Willing et al., 2018). IL-17+ MAIT are also associated with inflamed mucosal tissue, and are found activated and produce more IL-17 in IBD patients (Serriari et al., 2014). In addition to the gut, CD8+ MAIT cells have also been shown to be resident in normal skin and are thought to play a role in skin-associated inflammations, such as psoriasis and dermatitis herpetimorfis (Willing et al., 2018) and tissue repair (Constantinides et al., 2019). Therefore, it is tempting to speculate that 2DG or similar analogs of glucose could be used to modulate pathogenic responses of these T cell subsets.

An increase in IL-21 production in 2DG-treated memory T cells was also shown significant in our study. IL-21 has been documented to regulate the differentiation and function of several CD4+ T cells subsets, including Th17 cells (Leonard & Wan, 2016; Nurieva et al., 2007; Zhou et al., 2007), Th2 (Frohlich et al., 2007; Lajoie et al., 2014), and T follicular helper cells (TFH) (Vogelzang et al., 2008). Thus, the enhanced IL-17 and IL-4 cytokine production in 2DG treated T cells may be mediated partially by IL-21 signaling. IL-21 was also reported to induce IL-10 production in Th17 polarizing condition in mice (Spolski et al., 2009), which is consistent with our observation that differentiation of IL-17 secreting subsets from TNreg with 2DG increase IL-21 and IL-10 secretion. Furthermore, the functional significance of IL-21 in regulating effector functions of CD8+ T cells is highlighted by its critical role in sustaining anti-viral CD8+T cells during chronic LCMV infections (Cui et al., 2011; Elsaesser et al., 2009) and its potent effects to induce and expand cytotoxic CD8+ T cells for cancer immunotherapy (Davis et al., 2015; Santegoets et al., 2013). An increase production of IL-21 and IL-4 cytokines may suggest that 2DG enhance TFH cells, which have high expression level of CXCR5 and produce high level of IL-21 (Crotty, 2014). This hypothesis is also supported by that exogenous Ag-specific TFH cells do not require glycolysis (Choi et al., 2018). TFH is critical for the formation and maintenance of germinal centers and provide help for B cells generating antibody responses after immunizations (Bentebibel et al., 2013; Duan et al., 2014; Spolski & Leonard, 2010). Therefore, regulating TFH cells with glucose metabolism would potentially enhance development of new or improved vaccines (Crotty, 2014).

One potential mechanism of an increase frequency of IL-17-secreting T cell subsets in 2DG- treated cells is that 2DG alters T cells metabolism from glucose to alternative energy sources or metabolic pathways such as fatty acid oxidation or pentose phosphate pathway. Previous studies showed an inhibition of glucose metabolism with a glucose transporter inhibitor (CG-5) promoted fatty acid oxidation and the pentose phosphate pathway in CD4+ T cells in a similar manner as with 2DG (Li et al., 2019). Fatty acid metabolism has also been suggested to be involved in Th17 inflammation, as blockade of carnitine palmitoyltransferase 1 (CPT1), an enzyme responsible for catalyzing the breakdown of long-chain fatty acids, inhibits Th17-associated cytokine production from type 2 diabetes (T2D) patients (Nicholas et al., 2019).

One of our important findings towards mechanism of action of 2DG in our system is that 2DG- mediated increases in IL-17-secreting cells can be reversed by addition of equimolar mannose (Berthe et al., 2018; Kurtoglu et al., 2007). This suggests that another possible mechanism of 2DG is inhibiting the initial step of glycolysis affects the downstream metabolic pathway of mannose. Since mannose is a dominant monosaccharide in N-linked glycosylation (Imperiali & O’Connor, 1999), our observation may further suggest that part of the 2DG effects on T cell function are through modifying sugar moieties generated in central carbon metabolism (e.g. N-linked glycosylation). N-linked glycosylation plays a role in modulating activation and cytokine signaling (Baum & Cobb, 2017; Dean et al., 1979; Hauser et al., 2016), and thereby may affect differentiation and function of T cell subsets. An evidence to support this notion was our finding that mannose reversed 2DG-mediated down-regulation of CD25 expression upon activation. Similarly, another inhibitor of N-linked glycosylation, glucosamine, attenuated CD25 surface retention on T cells (Chien et al., 2015), and down-regulation of N-linked glycosylation promotes Th17 differentiation (Chien et al., 2015). However, surprisingly, addition of equimolar glucose could not reverse 2DG-mediated downregulation of CD25. We reason that 2DG efficiently inhibits glycolysis, therefore 2DG-treated cells could not utilize additional glucose. This speculation could be supported by seahorse glycolytic analysis showing that neither additional mannose nor glucose could reverse 2DG-mediated glycolytic capacity. Overall, it is conceivable that 2DG may have dual mechanism in enhancing the survival and effector functions of IL-17 secreting subsets.

Recently, 2DG has also been considered as a potential therapy during viral infections such as COVID-19 (Yang et al., 2021). Indeed, 2DG was approved by the Indian Council of Medical Research (OCMR) to treat COVID-19 patients in India (Sahu & Kumar, 2021). 2DG shortens the time of COVID patients recovered from the infection and there was a higher proportion of patients with improved symptoms who were free from supplemental oxygen dependence (Sahu & Kumar, 2021) . The underling mechanism of the effect of 2DG in these COVID patients is still unclear, but it was suggested that inhibiting glycolysis with 2DG could affected viral life cycle (Passalacqua et al., 2019) and an increased glucose level could enhance the suppressive function of monocytes and therefore enhanced SARS-CoV-2 infection (Codo et al., 2020). Given our findings, we also speculate that 2DG effects may have been through ameloriation of excessive or uncontrolled immune response during COVID19, which is observed during later severe disease stages. 2DG has also been tested to treat advanced cancer and proven safe in phase I studies (Raez et al., 2013; Stein et al., 2010). Our data shows long-term culture of human primary T cells with 2DG enriched IL-17 secreting T cells, which have been reported to exhibit pathogenic features (Basdeo et al., 2015). However, a significant increase in IFNγ production by 2DG may enhance therapeutic anti-tumor activity (Medrano et al., 2017). Taking together, our study provides insights for future clinical trials and strategies for development of 2DG-related cancer therapies.

## Materials and Methods

### T cell purification, activation, and culture

PBMCs from healthy individuals (New York Blood Center, New York, NY) were isolated using Ficoll-Paque plus (GE Healthcare). CD4+ T cells were isolated using Dynal CD4+ isolation kits (Invitrogen) and were 99% pure. Purified CD4+ cells were sorted in some experiments by flow cytometry (FACSAria; BD Biosciences) based on CD45RO, CCR7, CD25, and chemokine receptors expression into: 1) naive T cells (CD45RO-CCR7+CD25-), 2) memory T cells (CD45RO+CD25-), 3) naive regulatory T cells (Tregs; CD45RO-CD25+), 4) Th17 cells (CD45RO+CCR6+). Sorted subsets were 98% pure. All purified cells were kept at 37°C and 5% CO2 in complete RPMI 1640 media (RPMI 1640 supplemented with 10% FBS; Atlanta Biologicals, Lawrenceville, GA), 8% GlutaMAX (Life Technologies), 8% sodium pyruvate, 8% MEM vitamins, 8% MEM nonessential amino acid, and 1% penicillin/streptomycin (all from Corning Cellgro). To activate cells for expansion *in vitro*, anti-CD3/CD28 Dynabeads (Invitrogen) were used at a bead: cell ratio of 1:2 and the activated cells were cultured in complete RPMI 1640 medium (Thermo Fisher Scientific) or complete 1640 RMPI no glucose medium supplemented with IL-2 (10 ng/ml). For MAIT experiments, MAIT cells were activated by adding riboflavin metabolite 5ARU (Toronto Research, Ontario, Canada) into PBMC culture.

### Flow cytometry staining and analysis

Cells were stained with fluorochrome-conjugated Abs in FACS buffer (PBS+2% FCS and 0.1% sodium azide) for 30 min at 4°C (or room temperature for CCR6 staining). Abs used in surface staining are IL-2R (CD25), PD1, CD161, Va7.2, CCR6 (from BioLegend), LAG3, CD3, and CD8 (from Invitrogen). For intracellular cytokine staining, cells were stimulated for 4 h at 37°C with PMA (10ng/ml for PBMC and CD4+ T cells for 40 ng/ml) and ionomycin (500 ng/ml) (both from Sigma-Aldrich) together with GolgiStop (BD Biosciences). Cells were then stained with fixable viability dye (eBioscience) in PBS and surface marker Abs in PBS or FACS for 30 minutes, then fixed and permeabilized using eBioscience fixation/permeabilization buffers for 30 min at 4°C according to the manufacturer’s instructions, before staining for intracellular cytokines for 30 min at 4°C. Abs used for intracellular cytokines are IFNγ, TNF, IL-17A, IL-10, and IL-21(BioLegend). Ab used for Regulatory T cells transcription factor intracellular staining are Foxp3 and Helios Abs (BioLegend). Flow cytometry analyses were performed using an LSRFortessa X-20 flow cytometer (BD Biosciences) and SP6800 spectral cell analyzer (Sony Biotechnology).

### Seahorse metabolism assay

Seahorse assays were performed according to the manufacturer’s instructions to analyze glycolysis metabolism using the Seahorse XF Glycolysis Stress Test Kit (Agilent Technologies). Briefly, 14 day post activation, naive and memory T cells expanded in indicated conditions were washed and resuspended in glucose-free media (Gibco). 300k cells per well were spun down onto plates coated with Cell-Tak (Fisher scienctific). Four replicates were set up for each condition. Glucose, oligomycin, and 2DG were serially injected to measure metabolic function. Plates were analyzed using an XF^e^ 24 Extracellular Flux Analyzer (Agilent Technologies). Glycolysis was calculated as average post-glucose ECAR values minus average basal ECAR values.

### *In vitro* cytokine polarization assay

Sorted TN and TNreg were activated with anti-CD3/CD28 beads and cultured in complete RPMI media containing IL-2 10ng/ml (Chiron), together with IL-1β (10ng/ml), TGF-β (10ng/ml), and IL- 23 (100ng/ml) (R&D Systems). Cells were expanded for 2 weeks in media replenished for 2DG and IL-2.

### Statistical analysis

Data recorded by flow cytometry were analyzed using FlowJo (Tree Star, Ashland, OR). Statistical analyses were performed using GraphPad Prism 6.0 software (GraphPad Software, La Jolla, CA). Error bars represent SEM. Results were compared using two-tailed t tests. Bonferroni corrections were applied for multiple comparisons. For all experiments, significance was defined as *p < 0.05, **p < 0.01, ***p<0.001

## Acknowledgements

The research in this study was supported by National Institute of Health (NIH) grants U54NS105539 and R01AI121920 to D.U.

## Supplemental Figure Legends

**Supplemental Figure 1.**
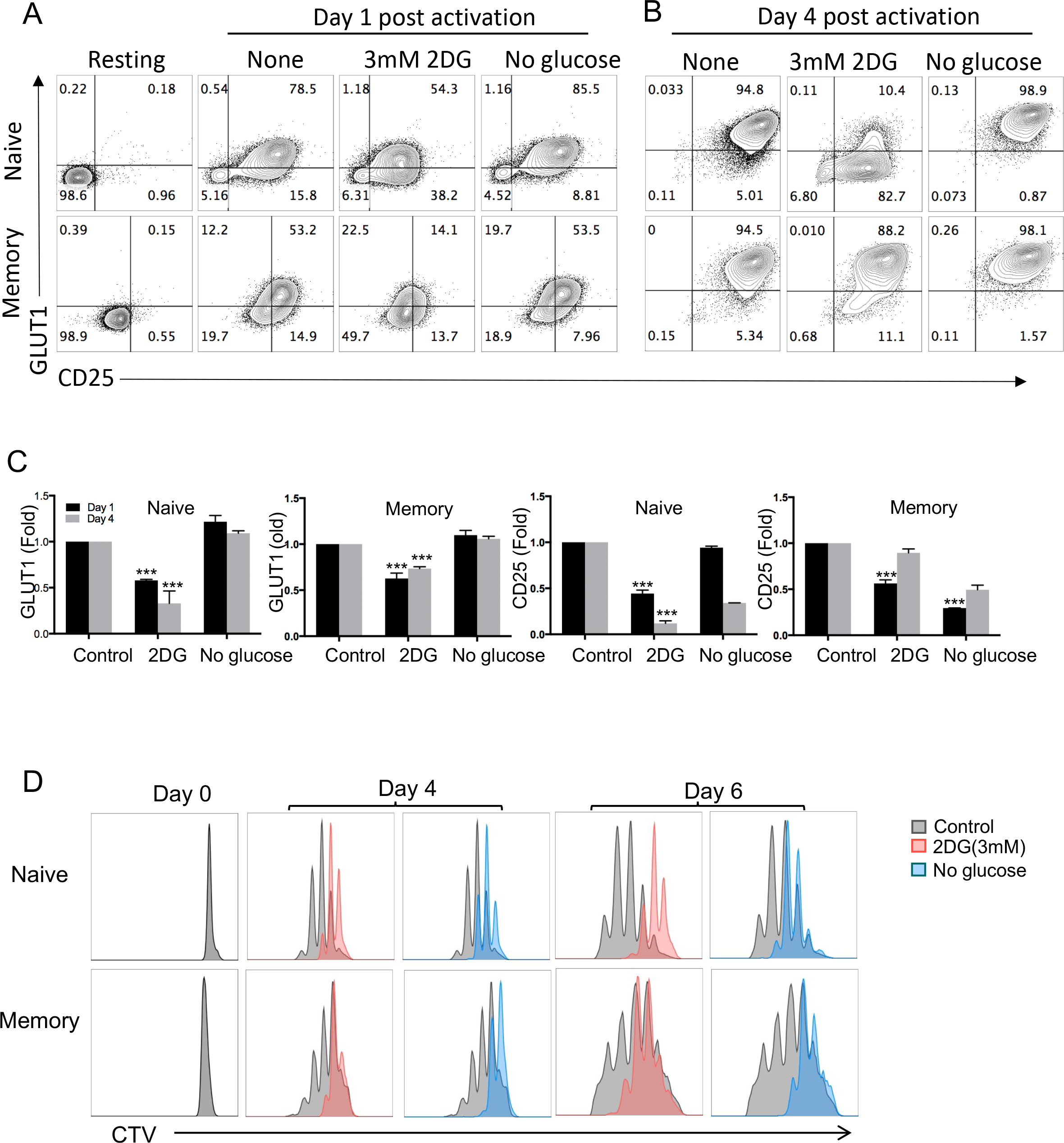
Addition of IL-2 did not restore 2DG effects on naive and memory T cell subsets. (A) Naive and memory CD4+ T cells were activated with aCD3/CD28 beads in media alone, 3mM 2DG, or in glucose-free media. Media was supplemented with IL-2. Representative flow cytometry plots of GLUT1 and CD25 surface expression on resting, and day 1-(A) or day 4- post(B) activated Naïve (top) and memory T cells (bottom) in indicated conditions were shown. (C) Fold difference in frequency of GLUT1 expression or mean fluorescence intensity of CD25 expression on naive or memory T cells on day 1- and day 4-post activation in media without IL-2. (D) Naive and memory CD4+ T cells were labeled with CTV dye followed by aCD3/CD28 beads activation in media alone (control), 3mM 2DG, or in glucose-free media. Representative histogram plot of CTV-labeled naïve (up) and memory (bottom) CD4+ T cells in 3mM 2DG (Red) or in glucose-free media(blue) after day 4 and day 6 post beads activation. Data represent three independent experiments. ***p<0.01

**Supplemental Figure 2.**
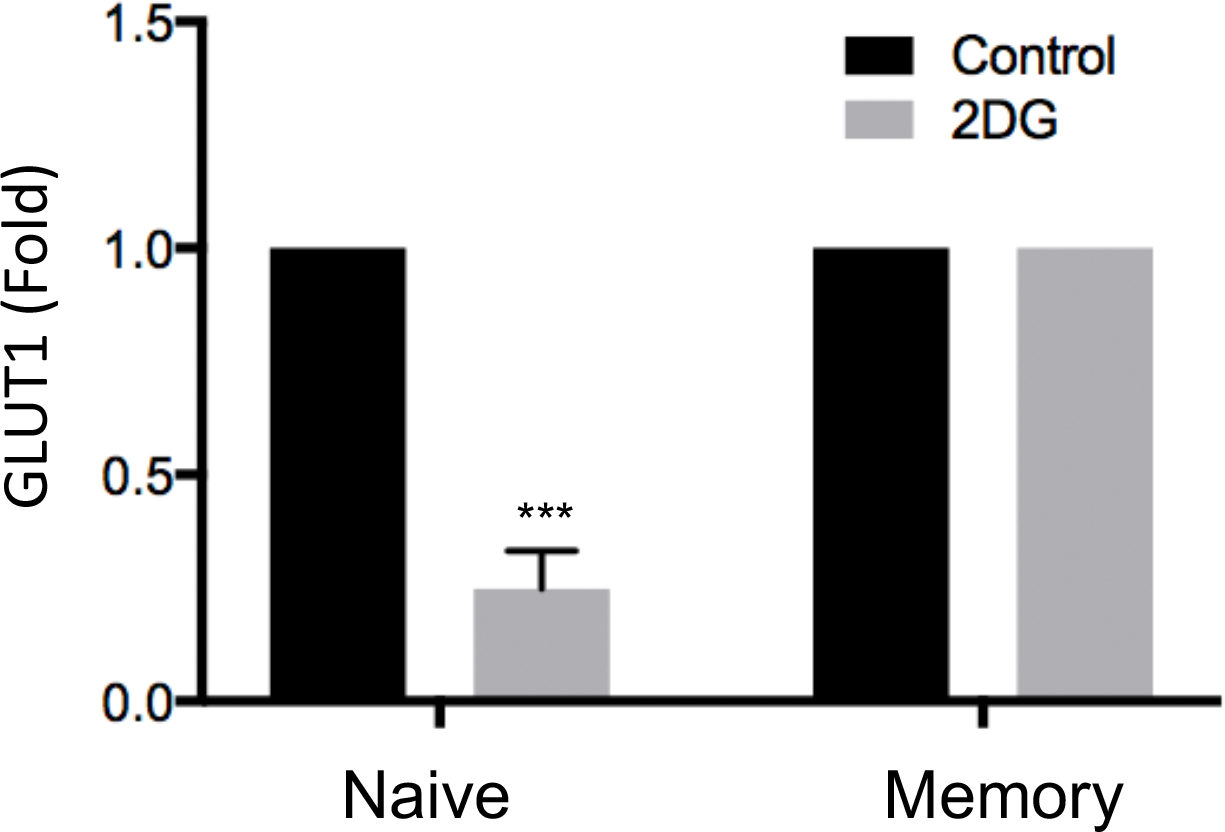
GLUT1 expression was down-regulated on naive T cells but not memory T cells after long-term culture. Naive and memory CD4+ T cells were activated with aCD3/CD28 beads in media alone (control in grey), or 1mM 2DG (in black). Media was not supplemented with IL-2. Fold difference in frequency of GLUT1 expression on naïve or memory T cells on day 14-post activation was shown. Data represent three independent experiments. ***p<0.01

**Supplemental Figure 3.**
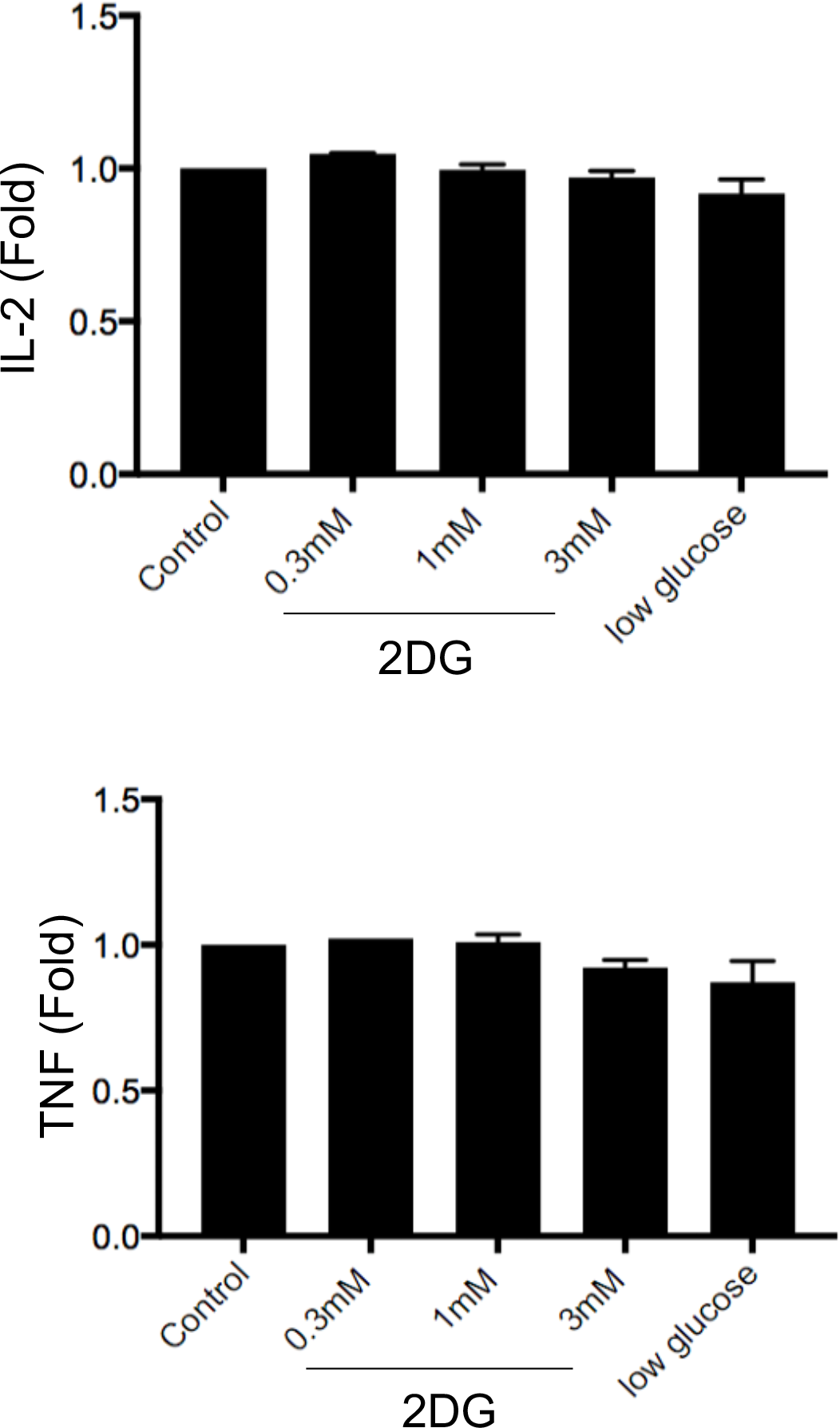
2DG did not change IL-2 and TNF production by CD4+ T cells. Purified CD4+ T cells were activated with aCD3/CD28 beads in media alone, with various doses of 2DG, and in glucose-free media. After 14 days’ cell culture, T cells activated and culture in indicated condition were re-stimulated with PMA/ionomycin for 4h, followed by intracellular staining as described in the method section. The percentage of IL-2 and TNF production from CD4+ T cells were shown. Data represent three independent experiments.

**Supplemental Figure 4.**
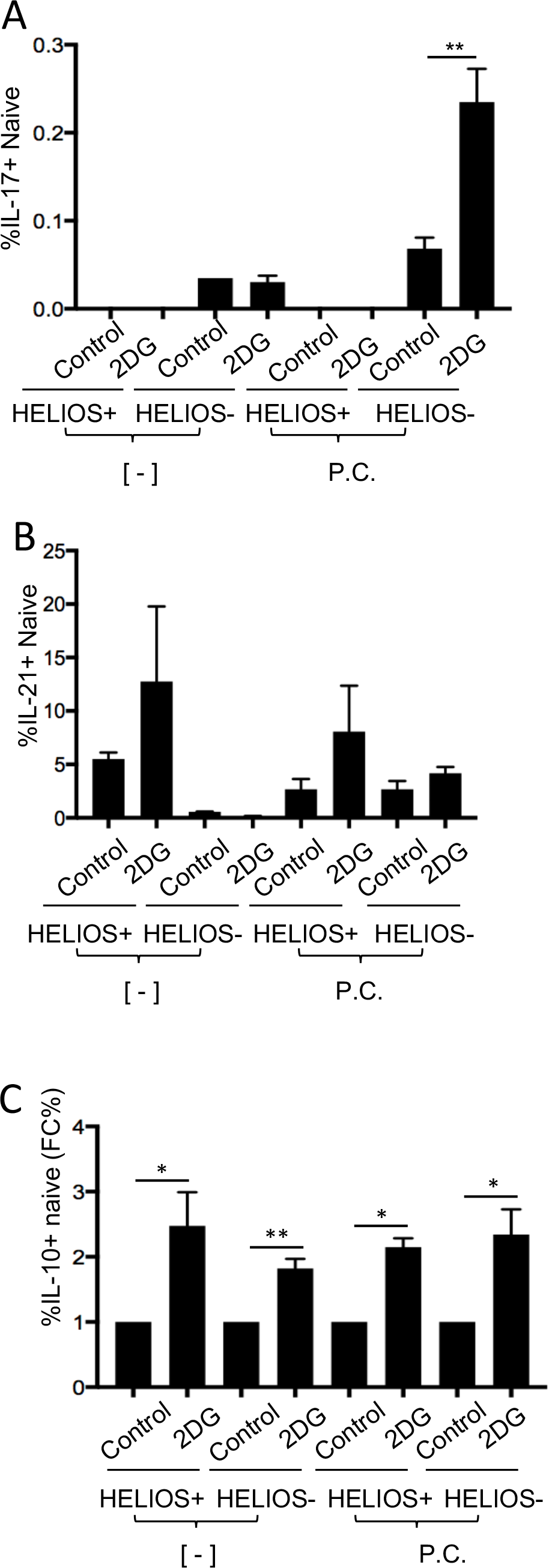
2DG enhances *in vitro* generation of IL-17-producing cells from naive T cells. Fold difference of IL-17 production from 1mM 2DG treated naive T cells with polarizing cytokines (P.C.) or without (-) compared with control after 14-day expansion.

**Supplemental Figure 5.**
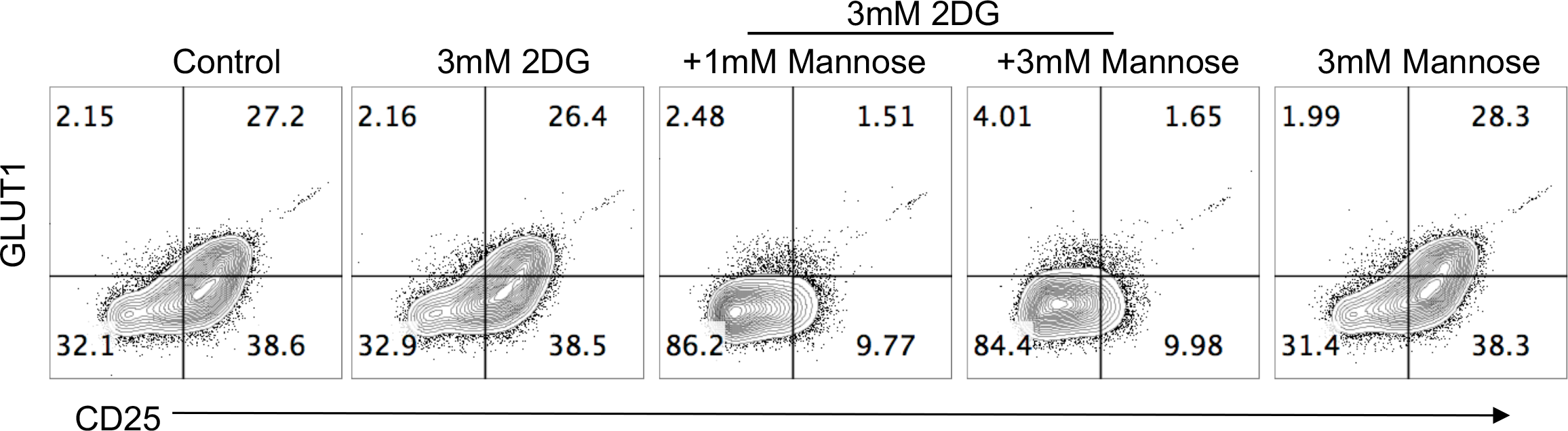
Mannose did not reverse GLUT1 expression 2DG-treated naive T cells. Naive CD4+ T cells were activated with aCD3/CD28 beads in media alone, 3mM 2DG alone, 3mM 2DG plus 1mM or 3mM mannose. Representative flow cytometry plots of GLUT1 and CD25 surface expression on naive T cells activated for 24 h in indicated conditions were shown. Data represent three independent experiments.

